# Nucleotide-dependent stiffness suggests role of interprotofilament bonds in microtubule assembly

**DOI:** 10.1101/098608

**Authors:** Katja M. Taute, Ernst-Ludwig Florin

**Author notes:** present address: The Rowland Institute at Harvard, 100 Edwin H Land Blvd, Cambridge, MA 02142, USA.

## Abstract

Many eukaryotic cell functions depend on dynamic instability, meaning the nucleotide-driven assembly and disassembly of microtubules. Assembly requires the constituent tubulin dimers to bind the nucleotide GTP, and its subsequent hydrolysis to GDP induces disassembly. The underlying structural mechanisms, however, are not well understood. Here, we determine the strength of contacts in the microtubule lattice by combining high precision measurements of the bending stiffness of analogues of GTP and GDP microtubules with a recent theoretical model. While previous structural studies have focussed on how the curvature of the tubulin dimer is affected by nucleotide binding, we present evidence of a dramatic regulation of the lateral interactions between the parallel protofilaments that dimers form in the microtubule. We conclude that the shear coupling between neighboring protofilaments is at least two orders of magnitude stronger in the GTP state than in the GDP state, and discuss the implications for the microtubule assembly.

## Introduction

Microtubules are essential to a wide variety of cell functions ranging from intracellular transport over directional polarity to cell division^1^. Many of these tasks critically depend on the microtubules’ ability to stochastically switch between growth and shrinkage in a process termed dynamic instability which is linked to the nucleotide state of the constituent protein subunits^2^. Dimers of *α* - and *β*-tubulin, which each need to bind a GTP (guanosine triphosphate) molecule, polymerize into a microtubule consisting of typically 13 parallel protofilaments. The GTP bound to the *β*-tubulin monomers is subsequently hydrolyzed to GDP (guanosine diphosphate), thereby rendering the microtubule lattice unstable and inducing depolymerization, unless a GTP cap on the growing end can be maintained by constant addition of subunits^3^.

The nature of the change in microtubule properties brought about by GTP hydrolysis has been the subject of much debate^4-6^. Recent efforts^7-9^have focussed on the intrinsic curvature of the tubulin dimer mediated by the longitudinal bonds within protofilaments as the source of dynamic instability. Less is known on the role of the weaker lateral contacts between protofilaments, and recent structural studies based on cryo-electron microscopy have yielded conflicting results. While some reports^10-12^support early conjectures^13,14^that these interprotofilament contacts are modified by the bound nucleotide, others^15,16^find no evidence of such changes. The fact that structural data^15^ do not reveal any major conformational differences between two microtubule states known to exhibit large differences in bending stiffness (stabilized with taxol versus polymerized with the GTP-analogue guanosine-5’-[(*α,β*)-methyleno]triphosphate, GMPCPP)^13,17-19^however suggests that current structural studies miss key features of microtubule internal contacts and highlights the need for alternative experimental techniques capable of resolving these.

Here, we exploit the dependence of the overall mechanical response of the microtubule on the contacts within the microtubule lattice to obtain information on the strength of interprotofilament coupling. Several previous studies have found evidence that the stiffness of microtubules is length dependent and attributed the effect to shear between the protofilaments softening the bending response of shorter microtubules^20-23^. Because the magnitude of the effect depends on the strength of the interactions between protofilaments, stiffness data can provide a readout of the interprotofilament contacts which are otherwise difficult to access experimentally. Comparing the length dependence of the bending stiffness of microtubules with different nucleotide contents, we employ a recent biopolymer model^24^ to show that the nucleotide has a strong effect on interprotofilament bonds and therefore likely also on microtubule assembly.

## Results

We obtain high precision stiffness data from the thermal fluctuations of grafted microtubules which are followed by tracking an attached fluorescent bead in two dimensions (Fig. 1 A-C)^21,23,25^. Mean square displacements (MSD) of the transverse coordinate are computed from the resulting time traces (Fig. 1D), and bending stiffnesses κ and first mode relaxation times τ_1_ are extracted under careful consideration of errors from sampling, correlations, low-pass filtering and position errors^25^ (see Methods). The resulting stiffness data are shown in Fig. 2 in terms of bending stiffnesses, κ, and equivalent persistence lengths, *l*_p_ = κ/(*k_B_T*).

**Figure 1.**
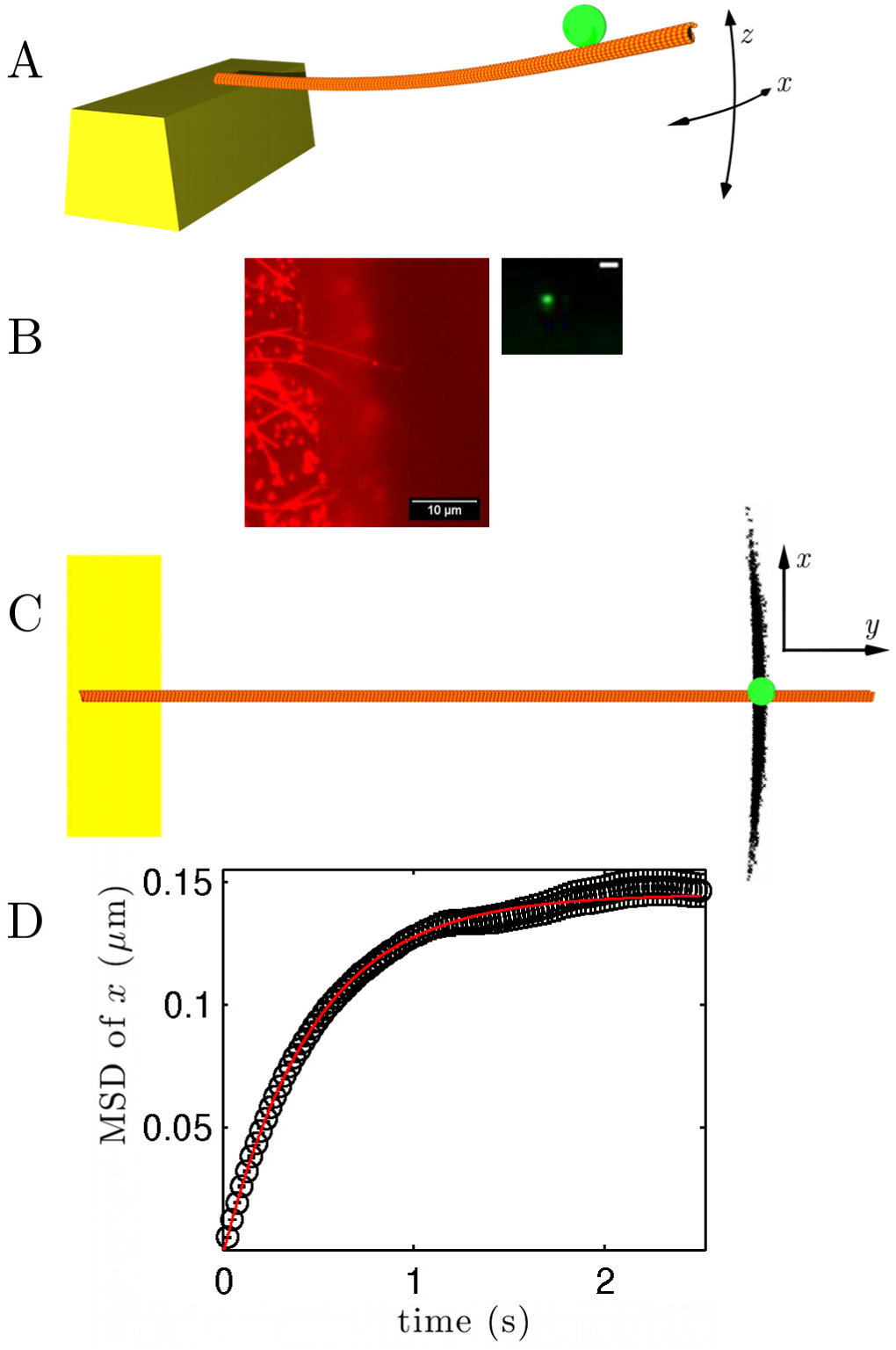
Schematic of setup. A) The free end of a grafted microtubule is subject to thermal fluctuations which are tracked by means of an attached fluorescent bead. B) Example fluorescence image of a rhodamine-labeled micrtobules (left) and the attached yellow-green fluorescent bead (right, scale bar 1 *μ* m). C) Imaging yields a two-dimensional projection of the bead’s position. Typically, the bead is tracked over 10,000-40,000 frames with a spatial precision of a few nm. D) Mean square displacement (MSD) of the transverse position (x) with fit of Eq. 10 for the same microtubule as in B and C.

### Taxol microtubules show a length-dependent stiffness and mechanical heterogeneity

Consistent with our previous study^23^, we find a positive correlation between microtubule length and stiffness for taxol microtubules (error-weighted Pearson’s correlation coefficient *r* = 0.46, *p* ≈ 2 × 10^-5^ under a two-tailed Student’s t-test). The scatter is larger than the error bars of the measurements, indicating heterogeneity in mechanical properties. For instance, for taxol microtubules with a length of ∼ 10 *μ*m, the measured stiffness values vary by up to a factor of 3. For microtubules shorter than ∼ 5 *μ*m, the stiffness appears approximately constant. Microtubules shorter than ∼ 2 *μ*m showed fluctuations too small and fast to be resolved, also indicating that their stiffness is not continuing to decrease any further.

### GMPCPP microtubules show a higher, but constant stiffness

Because the GMPCPP supports microtubule polymerization rates similar to GTP^26^, GMPCPP tubulin is widely considered an analogue of GTP tubulin^7,8,12,13,26^. GMPCPP is hydrolysed only slowly in the microtubule lattice and produces stable microtubules^26^.

We find that microtubules containing a high percentage of GMPCPP are approximately 3 - 10 times stiffer than taxol microtubules. In addition, however, our data reveal qualitative differences in mechanical properties. In contrast to the length-dependent stiffness of taxol microtubules, GMPCPP microtubules show a stiffness that is constant at *l*_p_ = 6400 ± 150 *μ*m (weighted mean ± unbiased standard error of the weighted mean) in the covered length range of 4.6 - 23.6*μ*m (*p* ≈ 0.997 for no correlation under a two-tailed t-test). This result is comparable to the value of *l*_p_ = 8800 ± 700μm found by Mickey & Howard^17^ for GMPCPP microtubules of lengths 28-64 *μ*m but covers a length regime that so far has not been probed rigorously (see Supplementary Discussion 1). Three separately prepared lots showed consistent results (Supplementary Fig. S1 and Supplementary Table S1 online). The thermal fluctuations of shorter GMPCPP microtubules are too fast to be reliably resolved with the CCD camera used in our assay.

The constant result for the stiffness of GMPCPP microtubules demonstrates that the scatter observed for taxol microtubules does not result from measurement error. Because taxol microtubules are softer, their fluctuations are slower and larger and hence less challenging to resolve. Therefore the associated measurement errors should be substantially smaller than for the stiffer GMPCPP microtubules.

### Taxol has little mechanical effect on microtubules

Given reports that taxol decreases^27^ or increases^17^ the stiffness of microtubules, measurements were also performed on GMPCPP microtubules with added taxol. The data show a small increase in stiffness compared to GMPCPP microtubules without taxol and more scatter (Fig. 2 and Supplementary Fig. S1 online). While the increase is statistically significant (*p* < 0.01 under a two-tailed Student’s t-test, see Supplementary Table S1 online), its magnitude corresponds to less than one standard deviation.

**Figure 2.**
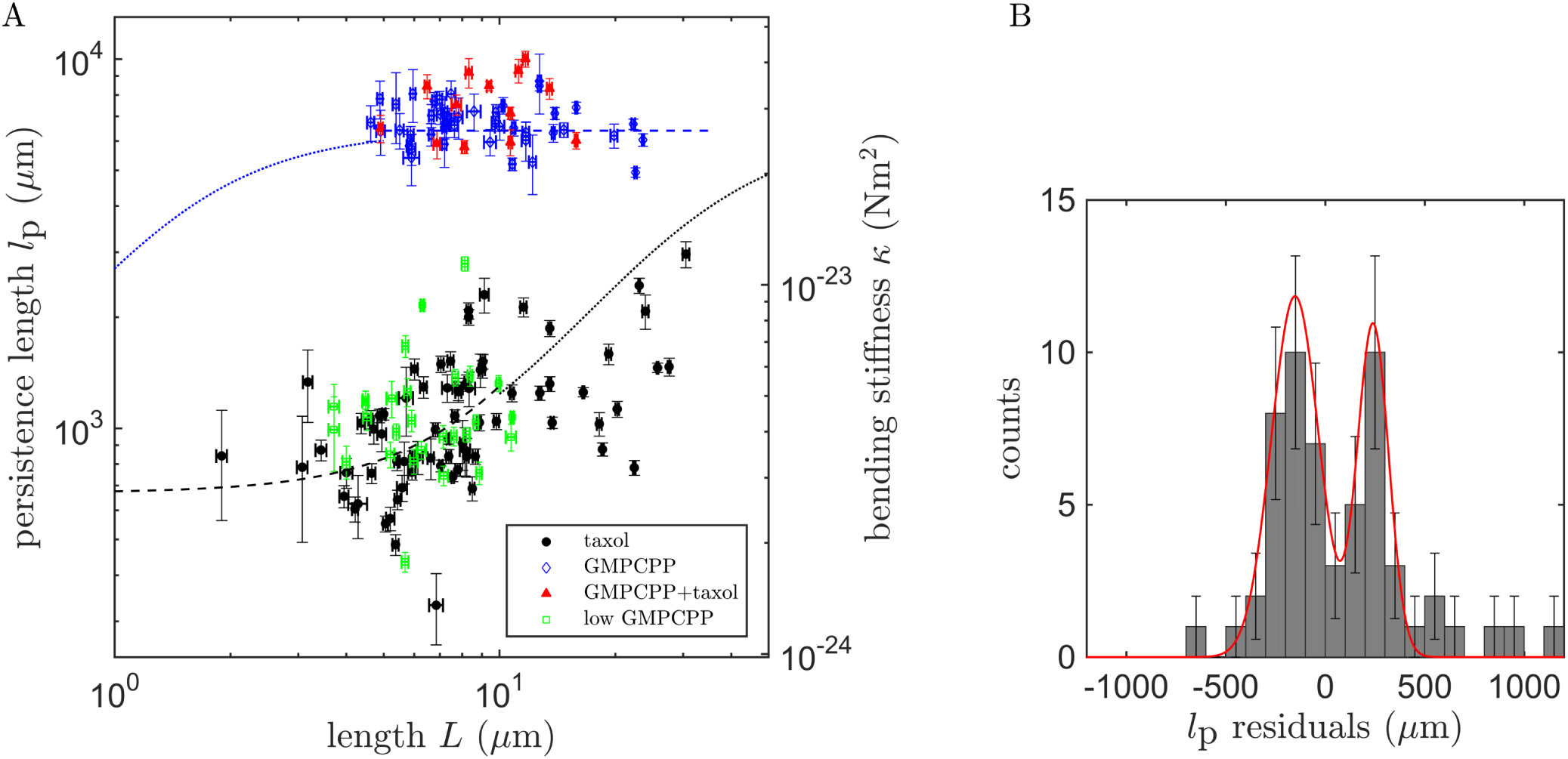
A) Bending stiffness measurements κ (right hand axis) and corresponding persistence lengths *l*_p_ = κ/(*k_B_T*) (left hand axis) for different microtubule types. The blue dashed line indicates the average value for GMPCPP microtubules. The black dashed line displays a fit of the Eq. 4 to the taxol microtubule data up to a length of 10 *μ*m. Dotted lines represent conjectures based on the WLB model. B) Histogram of residuals for the fit shown in A. Error bars represent Poisson noise, and the red line a fit of a sum of two Gaussians.

If the observed difference in stiffness between GMPCPP microtubules and taxol-stabilized GDP microtubules is a result of the nucleotide state, then one might expect a mixed nucleotide state to produce an intermediate effect. We therefore prepared microtubules that were polymerized with a mixture of GMPCPP and GTP (see Methods). As this procedure produced only a small number of microtubules that were stable for several hours, they likely contained a large fraction of GDP-bound dimers (see Supplementary Discussion 2). Surprisingly, the resulting stiffness data coincide with the ranges of values found for taxol microtubules and show a similar amount of scatter. This finding suggests that taxol also has little effect on the mechanical properties of GDP microtubules, and that taxol microtubules can hence be considered mechanical analogues of GDP microtubules.

### Relaxation times are consistent with stiffness data

Our analysis allows for the independent extraction of the bending stiffness, κ, and the first mode relaxation time, τ_1_ from the MSD. Because relaxation times depend on the stiffness, they provide a consistency check. Relaxation times are set by the interplay of stiffness and friction^28^:

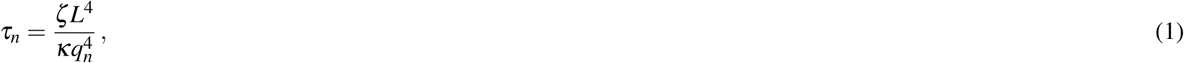

where L refers to the filament length, ζ to the friction coefficient per unit length, and *q_n_* to a dimensionless mode number. For the first mode of a grafted filament, *q*_1_ ≈ 1.875 ^29,30^.

Consistent with a constant stiffness, κ, we find that the relaxation times for GMPCPP microtubules follow an *L*^4^ scaling (see Fig. 3A). For taxol microtubules, however, the scaling is shallower, in agreement with our finding of a length-dependent stiffness κ(*L*). To explicitly determine whether stiffness and relaxation time values are consistent, ζ can be computed from them using Eq. 1. Fig 3B shows ζ (*L*) extracted for taxol microtubules. The fact that these values follow a clear trend and show very little scatter confirms that the scatter seen in the stiffness and relaxation time data is due to mechanical heterogeneity and not measurement error. The fit shown in Fig. 3B takes into account both hydrodynamic and internal friction (see Methods). Interestingly, the contributions from internal friction are similar in all types of microtubules investigated here, with a slight increase for GMPCPP microtubules (see Supplementary Fig. S2 & Table S2 online).

**Figure 3.**
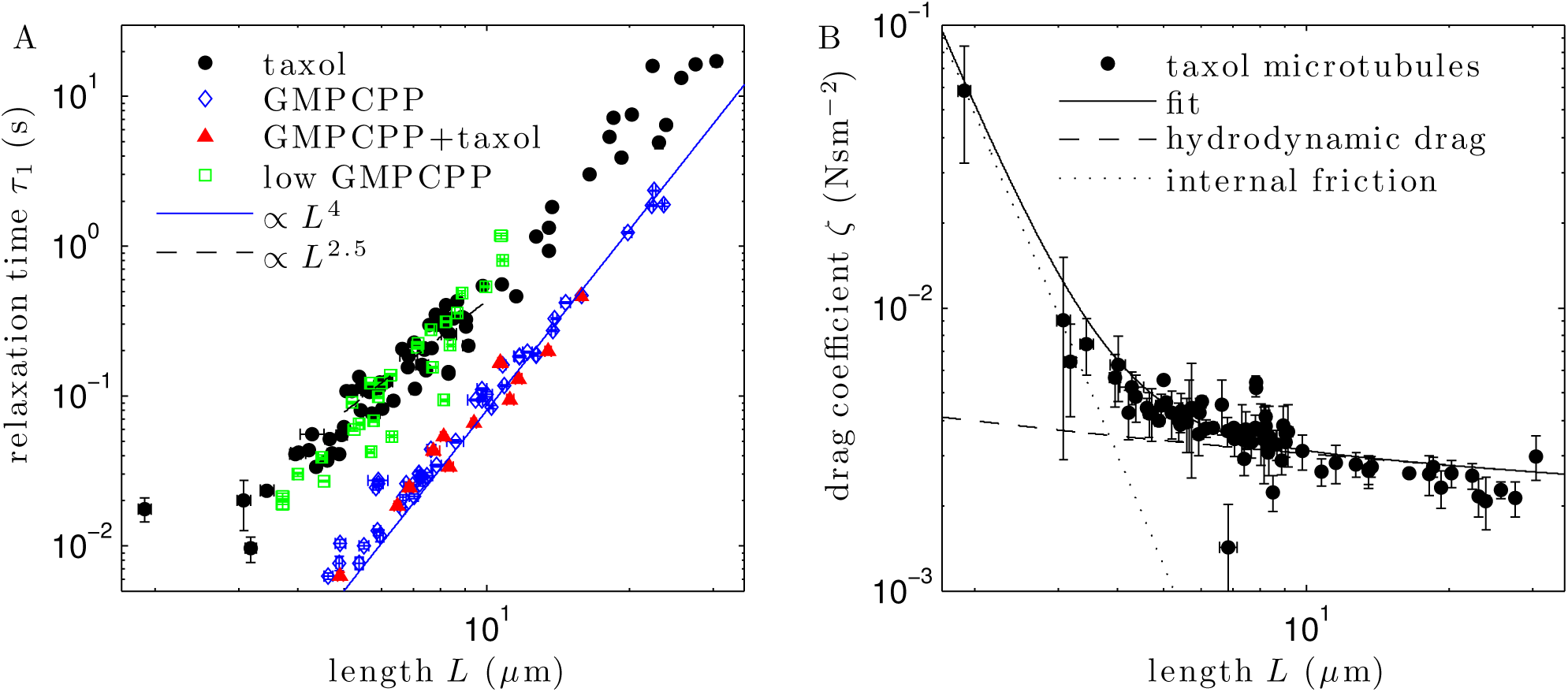
Dynamics confirm the stiffness data. A) First mode relaxation times, τ_1_, extracted from the MSD of the transverse coordinate. B) Drag coefficients per unit length, ζ, are computed from stiffness measurements and relaxation times using Eq. 1. In the interest of clarity, only data for taxol microtubules is shown. Data for other microtubule types are similar (Supplementary Fig. S2 online). The solid line represents a fit of Eq. 11 taking into account both hydrodynamic contributions and internal friction.

Knowing ζ(*L*), we can interpret the features of the relaxation time plots for taxol microtubules (Fig. 3A). The plateau around ∼ 20ms observed for the shortest taxol microtubules is the result of a sharp rise in ζ(*L*) due to internal friction. For microtubules longer than ∼ 5 *μ*m, Fig. 3B shows that ζ (*L*) is dominated by hydrodynamic effects and therefore approximately constant. Any deviation from an *L*^4^ scaling in the relaxation time data in this regime must therefore reflect a length dependence of the stiffness. A power law fit of the relaxation times for the regime 5 - 10*μ*m where the stiffness data display the clearest length dependence yields an exponent of 2.5 ± 0.2, similar to our previous finding of an approximate *L*^2^ scaling for microtubules shorter than 10 *μ*m^23^.

### Model of the length-dependent stiffness of taxol microtubules

In order to extract the strength of interprotofilament contacts, we employ the Wormlike Bundle (WLB) model developed by Heussinger et al.^24^ which is schematically depicted in Fig. 4. The WLB model considers the mechanics of a bundle of N weakly crosslinked protofilaments. The assembly is characterized by the effective force constants *k_x_* and *k_s_* resisting interprotofilament shear and protofilament extension, respectively, as well as the protofilament bending stiffness κ_f_ (see Fig. 4A-B). In the case of protofilaments arranged in a hollow ring as in a microtubule, Heussinger et. al.^31^ find the following expression for the resulting effective persistence length:

**Figure 4.**
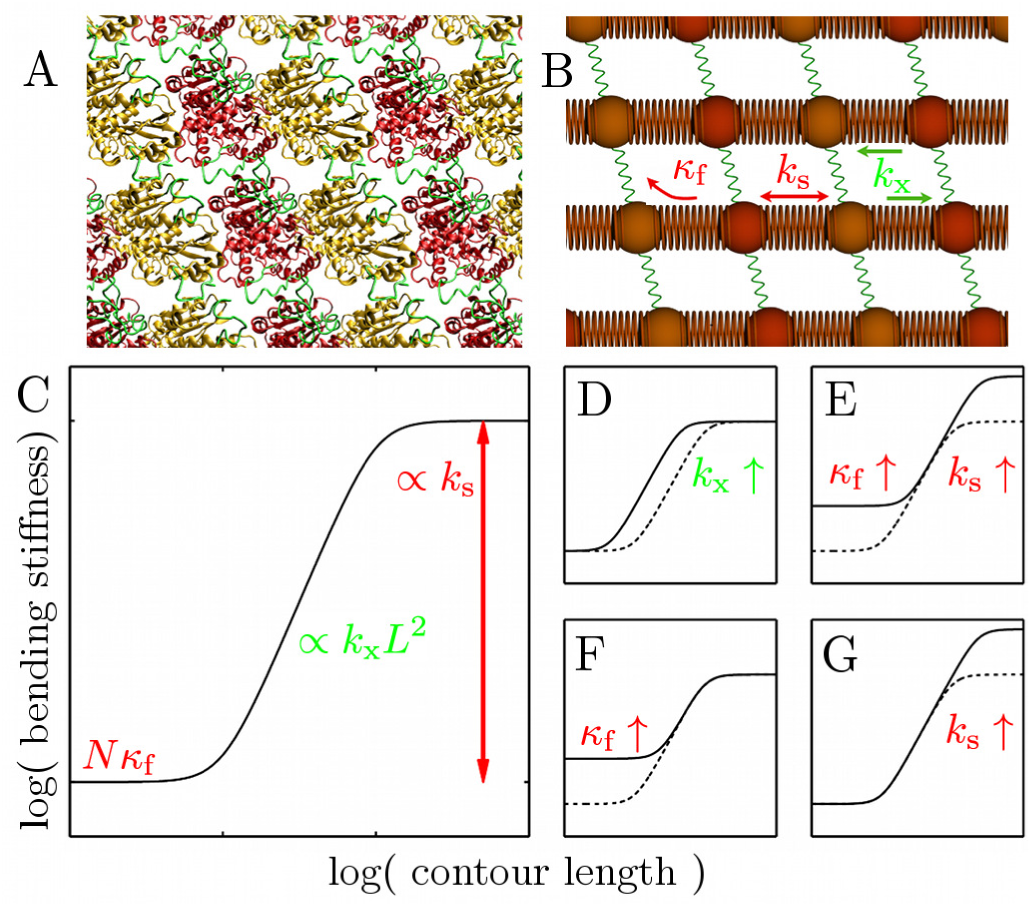
The WLB model applied to microtubules. A) A schematic of the microtubule lattice (based on PDB:1JFF^47^, created using VMD^48^) with the elements thought to be involved in interprotofilament contacts^49^ highlighted in green. B) WLB model of the microtubule lattice. Monomers (shown as spheres) are connected by strong longitudinal interactions (red springs with force constant *k_s_)* along the protofilaments. Neighbouring protofilaments are coupled laterally with soft shear springs (green springs with force constant k_x_). Each protofilament also has a bending stiffness K_f_. C) Schematic of the resulting bending stiffness as a function of length as predicted by the WLB model. Three distinct scaling regimes are each dependent predominantly on one of the three parameters. D) Schematic of how changes in those parameters affect the resulting stiffness.

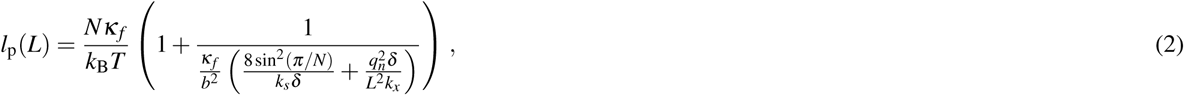

where δ and *b* refer to the longitudinal spacing of crosslinks and the lateral spacing of protofilaments, respectively. Figs. 4C-D illustrate the three distinct scaling regimes of this expression, each of which depends almost exclusively on only one of the three parameters. For very long microtubules, the assembly behaves as though fully coupled and shows a constant persistence length

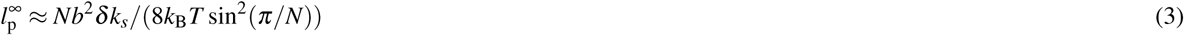

determined by the extensibility of the protofilaments, analogous to the Young’s modulus in beam mechanics. For very short microtubules, the stiffness is given by the sum of the bending stiffnesses of the individual protofilaments, 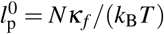. In the intermediate regime, the stiffness of the assembly is governed by interprotofilament shear, resulting in a scaling proportional to *k_x_L^2^*.

Our data for taxol microtubules support a constant stiffness for taxol microtubules shorter than ∼ 5 *μ*m, and an increasing stiffness for longer ones. A stiffness plateau of 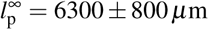 was previously found by Pampaloni et al.^21^ for taxol microtubules longer than ∼ 40 *μ*m. Our present measurements only cover lengths up to ∼ 30 *μ*m and do not include this plateau. Therefore, we employ a simplified version of Eq. 2 valid far below 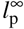:

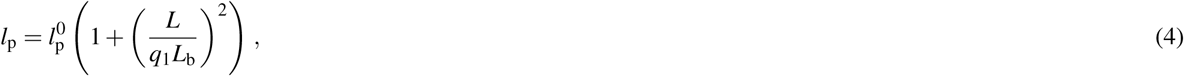

where

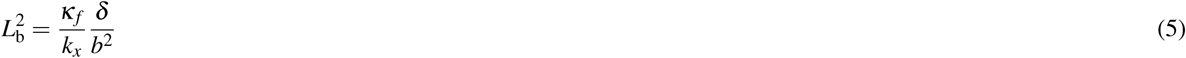

marks the characteristic length scale of the transition between constant and increasing stiffness. A fit of the data up to lengths of 10*μ*m yields 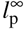 and *L_b_* = 5.5 ± 0.2*μ*m and is shown as a dashed black line in Fig. 2. The dotted continuation shows the full version of Eq. 2 where 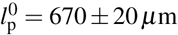 is assumed to equal the mean stiffness of GMPCPP microtubules, *l*_p_ = 6400 ± 60 μm,which, strinkingly, coincides well with the value 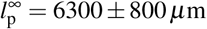 Pampaloni et al.^21^ found for long taxol microtubules. The constant stiffness we find for GMPCPP microtubules may therefore correspond to the same limit 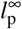. If so, then the extensibility *k_s_* of protofilaments is unchanged as a result of the different nucleotide. With Eq. 3 and the parameters δ ≈ 4nm^4^ and *b* ≈ 5nm^32^, we obtain *k_s_* ≈ 7N/m which is consistent with the longitudinal position fluctuations of taxol-bound tubulin dimers in molecular dynamics simulations^33^.

### Mechanical heterogeneity of taxol microtubules

The large scatter we observe for taxol microtubules suggests underlying subpopulations with different mechanical properties. The source of this mechanical heterogeneity likely lies in variations in microtubule architecture. While in vivo, the protofilament number is tightly controlled to 13 in most cases, in vitro numbers from 9 to 16 have been observed^34^. For GTP-induced polymerization followed by addition of taxol, it has been reported that 13, 14 and 15 protofilaments dominate the distribution which remains stable over the course of days^35^. The residuals of our fit of Eq. 4 (Fig. 2B) provide evidence for two mechanically distinct subpopulations.

Can protofilament variation quantitatively account for the scatter in measured stiffnesses? If the microtubule is modeled as a cylinder made from a homogenous, isotropic material, a change in protofilament number from 13 to 14 should only result in a modest change in stiffness of less than 20%^36^, while experimentally a variability by a factor of 3 is observed for microtubules of similar lengths.

The WLB model, however, does provide a mechanism by which differences in protofilament number could produce a large difference in stiffness. It is known from structural data that, for microtubules with protofilament numbers other than 13, the lattice shows a supertwist around the microtubule axis with a pitch on the order of several micrometers^32,37^. Such helicity is expected to drastically lower the stiffness in the WLB bundle model. Because protofilaments change side going along the length of a bent microtubule, they are able to accommodate the deformation more easily^31^. This softening can only affect the intermediate and long length regime. As a result, only 13 protofilament microtubules are expected to follow the idealized WLB scaling depicted in Fig. 4C.

In addition, lattice defects^38^ may contribute further mechanical heterogeneity for long microtubules.

### Nucleotide-dependent lateral interactions between protofilaments

If the constant stiffness observed for GMPCPP microtubules corresponds to the limit 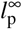 then the length-dependent regime expected from shear contributions must have been shifted to shorter microtubule lengths, corresponding to a strong increase in the interprotofilament shear constant *k_x_* (compare Fig. 4D). In the length regime covered by our measurements, we see no indication of a decrease in stiffness for shorter microtubules. In order to obtain a lower bound estimate for the shear force constant *k_x_*, we therefore assume that for the shortest GMPCPP microtubule measured, any shear-induced decrease in stiffness from 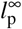 cannot be larger than the measurement errors. This microtubule had a length of *L*_1_ = 4.6 ± 0.1 *μ*m and a persistence length of 6730 ± 740*μ*m, and we approximate the error bars as 10%. Eq. 2 with the simplification that 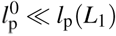 yields a shear force constant of 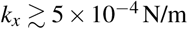 per monomer.

For taxol microtubules, by contrast, Eq. 5 together with our fitted values for *L_b_* and 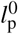 provide us with an estimate of *k_x_* ≈ 10^-6^ N/m. A change by more than two orders of magnitude in the lateral interaction between protofilaments may seem extreme at first glance, but may simply reflect the similarly extreme differences in polymerization behaviour. While GMPCPP tubulin readily polymerizes into stable microtubules^26^, GDP tubulin has a low affinity for the microtubule lattice and, once incorporated, promotes depolymerization^2^.

## Discussion

Our results demonstrate for the first time that the bound nucleotide causes not just a quantitative change in microtubule stiffness but also a qualitative change in the mechanical behaviour of microtubules, resulting in a length-dependent versus a constant stiffness for GDP versus GMPCPP microtubules, respectively, in the length regime of approximately 5 - 24 *μ*m covered here.

Our interpretation of a nucleotide-dependent interprotofilament contact strengths may have strong implications for microtubule assembly. An increased shear coupling in the GTP state would likely aid polymerization by locking tubulin dimers into the correct geometry. Interestingly, GMPCPP microtubules show both a higher nucleation rate and a higher homogeneity in the resulting microtubule architecture than microtubules polymerized with GTP, but their polymerization rate is almost identical^4,26^. While GMPCPP microtubules have been shown to almost exclusively consist of 14 protofilaments, microtubules polymerized from GTP tubulin in vitro typically have protofilament numbers varying from 12 to 15, with 13 and 14 protofilaments being the most common^4^. The discrepancies raise the question whether GMPCPP tubulin really is a good mimic of GTP tubulin as is commonly assumed.

One explanation that could reconcile this assumption with the observations is that for GTP tubulin, hydrolysis or intermediate steps already begin soon after the assembly of dimers into the microtubule lattice. In the face of a weak lateral coupling for the protofilaments in the hydrolysed part of the microtubule, strong lateral contacts in a small GTP cap might not be sufficient to prevent stochastic longitudinal sliding, leading to variations in protofilament architecture and lattice defects. The weak lateral coupling in the GDP state might also explain why protofilaments can take on such a vast variety of relative geometries, from different microtubule protofilament numbers and helix rises over antiparallel sheets^39^ to even comma-shaped arrangements^40^. For GMPCPP microtubules, however, the strong lateral coupling may prevent longitudinal sliding and lock the microtubule lattice into one particular geometry. The recent finding that the size of the GTP cap is several times larger in vivo than in vitro^41^ might hence be related to the more tightly controlled microtubule architecture in vivo.

Our interpretation of microtubule mechanics based on the Wormlike Bundle Model accounts for all the features observed in our mechanical data. Its particular strength lies in providing a unified explanation of how microtubule architecture affects stiffness to cause disparate effects such as the length dependence and scatter observed for taxol microtubules as well as the constant stiffness observed for GMPCPP microtubules. Some of its quantitative implications are however challenging to reconcile with the lattice rotation model^42^. To explain why, for protofilament numbers other than 13, the microtubule lattice twists around the microtubule axis, it was proposed that the lateral contacts between protofilaments are resisting a longitudinal mismatch created by the changed number of protofilaments. It is unclear at present how a weak lateral coupling between GDP protofilaments might create the force necessary to maintain a supertwisted microtubule lattice. Further quantitative work will be necessary to resolve this conflict.

In conclusion, we have shown that the bound nucleotide modulates microtubule mechanics both in a quantitative and in a qualitative fashion, changing not only the mean stiffness but also inducing a length dependence for GDP microtubules which is not observed for a GTP analogue. The observed data are consistent with an interpretation of microtubule mechanics based on interprotofilament shear. Our interpretation suggests that strong interprotofilament contacts in the GTP state assist microtubule assembly, and that nucleotide hydrolysis alters microtubule mechanics by weakening the lateral contacts between protofilaments.

## Methods

### Microtubule polymerization

Taxol microtubules were polymerized at 37° C and ∼4mg/ml tubulin with 1 mM GTP in a polymerization buffer consisting of BRB80 (80mM PIPES, 1 mM EGTA, 1 mM MgCl_2_, pH 6.8) supplemented with additional 4mM MgCl_2_. High GMPCPP microtubules were prepared from tubulin which had been cleared from other nucleotides by a spin column buffer exchange followed by a polymerization-depolymerization step, and assembled overnight at 0.5-1mM GMPCPP, 0.1-0.4mg/ml tubulin, and temperatures between 23 and 30°C. Low GMPCPP microtubules were polymerized for 3h at 33°C using 0.8mg/ml tubulin, 1 mM GMPCPP, and ∼0.1 mM GTP. After polymerization, all microtubule solutions were cleared of free tubulin by centrifugation and resuspension in BRB80 (supplemented with 20μM taxol in the case of taxol microtubules). In all cases, an approximate ratio of 10:2:1 was used of unlabeled, rhodamine-labeled and biotinilated tubulin (Cytoskeleton, Denver, CO). Taxol microtubules were used for up to 5 days after polymerization, high GMPCPP microtubules up to one day after polymerization, and low GMPCPP microtubules were always used freshly.

### Sample preparation

Samples were prepared as described previously^21,23,25^. Clean TEM gold grids were glued onto a round 15mm coverslip with silicone glue (Elastosil N10, Wacker Chemie, Germany) and, after drying, cleaned by submerging in a solution of 5% hydrogen peroxide and 5% ammonia in water for 15 min. After extensive rinsing with water and then with ethanol, the grids were functionalized by submerging them overnight in a 5mM solution of 11-mercaptoundecanoic acid in ethanol. After extensive rinsing with ethanol, the grids were dried with nitrogen and assembly into a flow chamber consisting of a 24x32mm #1 coverslip (Menzel Gläser, Germany), two strips of parafilm and the round coverslip on top with the grid facing inward. Brief heating on a hot plate sealed the parafilm to the glass, and solutions could then be exchanged in the ∼ 10 *μ*l volume by blotting with tissue. The grids were then activated by flowing in freshly prepared 100mM N-hydroxysuccinimide and 100mM N-(3-dimethylaminopropyl)-N-ethylcarbodiimide hydrochloride in BRB80. After 20 min, the chambers were flushed with 500*μ*l of BRB80 before microtubules were flushed in and allowed to bind to the gold surfaces for 30-60min, then 200nm, yellow-green, neutravidine-coated beads (F-8774, Invitrogen, Carlsbad, CA) were flushed in and left to bind to the biotinilated microtubules for 20min. To remove unbound beads and microtubules, the sample was then flushed with 200*μ*l of BRB80 containing an oxygen scavenger system^25^ of ∼ 0.3mg/ml glucose oxidase, ∼ 0.3mg/ml catalase, 30mM glucose, and 0.3% saturated hemoglobin, and sealed with valap. For taxol microtubules, the buffer solution also contained 20 *μ*M taxol.

### Microscopy

On an inverted fluorescence microscope (Axiovert 10, Zeiss) modified for increased mechanical stability and equipped with a water immersion objective lens (UPlanSApo60xW, NA 1.2, Olympus) the samples were visually inspected and microtubules were selected that showed clear attachment to the gold substrate, no overlap with other microtubules, and a bead attached somewhere along their free length. Using a CCD camera (Sensicam QE, PCO), 100-300 frames were recorded using a rhodamine filterset to facilitate microtubule length measurements. Then, 1 - 4 × 10^4^ frames were recorded of the yellow-green bead without further fluorescence excitation of the microtubule. Integration times and frame rates were adjusted in the range of 14-80 ms and 10-70 Hz, respectively, to adequately capture the first mode fluctuation dynamics. Images were recorded using PCO Camware and exported as 16-bit TIFF files. All measurements were carried out at room temperature (21-23°C).

### Theory

The theoretical expression for the transverse mean square displacement (MSD) at a position s along a thermally fluctuating grafted filament of length *L* is

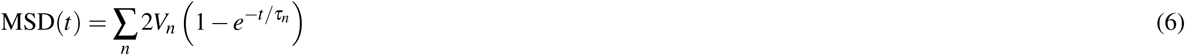

where

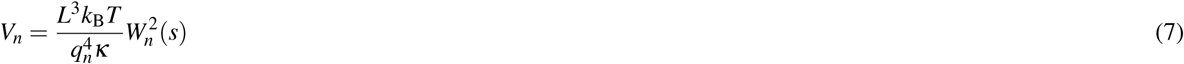

is the position variance due to the *n*^th^ mode. The term *W_n_(s)* refers to the contribution of the *n*^th^ eigenmode at position *s*, and is given by^30^

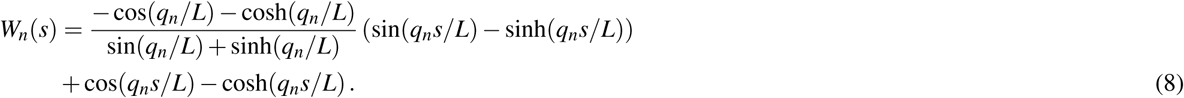

The mode numbers *q_n_* are given by *q_1_* ≈ 1.875, *q*_2_ ≈ 4.694, *q*_3_ ≈ 7.855, and *q_n_* ≈ (*n* - 1/2)π for higher modes. Modes higher than the first are typically not resolved and therefore only contribute an offset *a* to Eq. 6, reducing the expression to

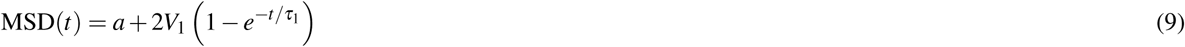

Any errors in the position determination will also contribute an offset to the MSD and can therefore be absorbed into the fit parameter *a*. The first mode can be subject to low-pass filtering because the motion is blurred if the integration time *T* is comparable to the relaxation time τ_1_. The expression^23^

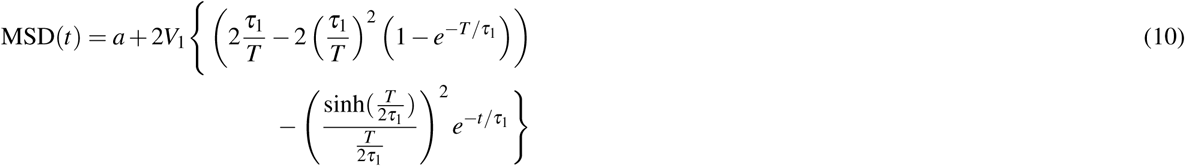

accounts for this effect and was used to fit the experimental data. Fit parameters are the offset *a,* the first mode variance *V*_1_, and the first mode relaxation time τ_1_. Using Eq. 7, the stiffness κ can then be extracted from *V*_1_.

### Data analysis

Microtubules lengths were measured using the line segment measuring tool in ImageJ^43^ on frames where the microtubules showed little curvature. Particle tracking and data analysis were carried out using custom-written Matlab code. We obtained data from 26 bovine and 26 porcine taxol microtubules which we combined with previous measurements^23^ of 20 bovine taxol microtubules as no systematic differences between porcine and bovine tubulin were apparent (Supplementary Fig. S3 online). For consistency, all data were analyzed using the same methodology. We obtained 44 measurements on 41 GMPCPP microtubules, 28 measurements on 27 low GMPCPP microtubules, and 13 measurements on 12 GMPCPP taxol microtubules. Microtubules with two beads attached at different positions along their length allowed for two independent measurements.

The MSD was computed for the transverse position coordinate and error estimates were obtained using the blocking procedure^44^. The data were then fit by Eq. 10. The fits were weighted by error estimates. The shortest microtubule included in our previous study^23^ had to be excluded because the larger number of fit parameters used in the present stricter analysis procedure led to unacceptably large error bars.

The fits shown in Figs. 2 & 3 were performed in logarithmic space and weighted by relative error. Drag coefficients are fit by the equation

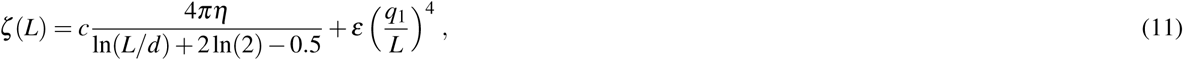

where the first and second term refer to contributions from hydrodynamic^45^ and internal friction^46^, respectively. The parameter *d* gives the diameter of the microtubule and η the viscosity of the medium, and *c* and ε are fit parameters.

## Acknowledgements

The authors would like to thank Prof. Tim Mitchison, Prof. Ken Johnson, Dr. Claus Heussinger and Prof. Joe Howard for stimulating discussions, and Prof. Marileen Dogterom for helpful comments on the manuscript. K.M.T. gratefully acknowledges support by a doctoral fellowship from the Gottlieb Daimler and Karl Benz Foundation. This research was funded by NSF grants CMMI-0728166 and CMMI-1031106.

## Author contributions statement

K.M.T. and E.-L.F. conceived the experiments, K.M.T. conducted the experiments and analyzed the data, K.M.T. and E.-L.F. interpreted the results and wrote the manuscript.

## Additional information

**Competing financial interests** The authors declare no competing financial interests.

